# Tiger team: a panel of human neutralizing mAbs targeting SARS-CoV-2 spike at multiple epitopes

**DOI:** 10.1101/2020.05.20.106609

**Authors:** Tal Noy-Porat, Efi Makdasi, Ron Alcalay, Adva Mechaly, Yinon Levy, Adi Bercovich-Kinori, Ayelet Zauberman, Hadas Tamir, Yfat Yahalom-Ronen, Ma’ayan Israeli, Eyal Epstein, Hagit Achdout, Sharon Melamed, Theodor Chitlaru, Shay Weiss, Eldar Peretz, Osnat Rosen, Nir Paran, Shmuel Yitzhaki, Shmuel C. Shapira, Tomer Israely, Ohad Mazor, Ronit Rosenfeld

## Abstract

The novel highly transmissible human coronavirus SARS-CoV-2 is the causative agent of the COVID-19 pandemic. Thus far, there is no approved therapeutic drug, specifically targeting this emerging virus. Here we report the isolation and characterization of a panel of human neutralizing monoclonal antibodies targeting the SARS-CoV-2 receptor binding domain (RBD). These antibodies were selected from a phage display library constructed using peripheral circulatory lymphocytes collected from patients at the acute phase of the disease. These neutralizing antibodies are shown to recognize distinct epitopes on the viral spike RBD, therefore they represent a promising basis for the design of efficient combined post-exposure therapy for SARS-CoV-2 infection.

## Introduction

The present global pandemic of coronavirus induced disease 19 (COVID-19), declared by the World Health Organization (WHO) as a public health emergency of international concern, is caused by the highly transmissible severe acute respiratory syndrome coronavirus 2 (SARS-CoV-2). To this date, about 5 million confirmed cases and over 300,000 deaths have been reported worldwide^1^. Yet, there is no approved therapeutic drug specifically targeting the SARS-CoV-2. The novel coronavirus SARS-CoV-2, emerged as the seventh type of coronavirus infecting humans and the third most pathogenic preceded by the Severe Acute Respiratory Syndrome coronavirus (SARS-CoV) and the Middle East Respiratory Syndrome coronavirus (MERS-CoV).

Due to their exceptional antigen specificity, therapeutic monoclonal antibodies (mAbs) are considered an attractive candidate to target exposed antigenic sites on viruses, and prevent their infectivity^2^. The therapeutic efficacy of mAbs, specifically targeting surface viral proteins was demonstrated before for several viruses including SARS-CoV, MERS and Ebola^3-6^.

SARS-CoV-2 utilizes the surface homotrimeric Spike glycoprotein (S) as a major mediator of cellular infection. The SARS-CoV-2 S protein is composed of two distinct subunits, S1 and S2. The S1 subunit contains the receptor binding domain (RBD), known to bind the Angiotensin-Converting Enzyme 2 (ACE2) receptor on host cell surfaces^7^. The S2 subunit mediates the fusion of the viral and cellular membranes, essential for viral entry into the cell. The receptor interaction site on S1 is considered the main target for efficient neutralization of cell infection and therefore a prime candidate for therapeutic antibody development^8-12^. Although the S protein of the SARS-CoV and SARS-CoV-2 share 77.5% identity, most of the mAbs isolated against SARS-CoV reportedly failed to cross-neutralize the novel virus^13-15^.

For designing optimal therapeutic strategies, there is an urgent need for the identification of neutralizing monoclonal antibodies that specifically target SARS-CoV-2 (such antibodies may be identified either in humans in the course of illness/recovery, or in immunized animals). Recently, first two mAbs, elicited against the SARS-CoV-2, were reported^16^. Furthermore, efficient post-exposure therapy in humans, may require integration of several noncompeting mAbs, ideally neutralizing the virus infectivity by different mechanisms. Such combined therapies are expected to provide superior control of potential neutralizing escape variants^9,17^.

Here we describe the isolation of a panel of neutralizing mAbs selected against the SARS-CoV-2 RBD from phage display library constructed based on patient samples collected in the acute phase of the disease. These specific antibodies were found to recognize distinct epitopes and can potentially be used either for therapy or immune prophylaxis.

## Materials and methods

### Blood samples

Sera and whole blood samples collected from five convalescent or severe COVID-19 patients were obtained under written inform consent and treated in accordance with the biosafety guidelines of the IIBR in BL3 facility. PBMCs were separated from fresh whole blood using density centrifugation by Ficoll. Sera samples were heat-inactivated (30 min at 60°C) prior to use for binding or neutralizing assays. The study was approved by the Sheba Medical Center IRB Ethical Committee as well as by the Baruch Padeh Medical Center IRB Ethical Committee.

### Phage Display scFv libraries construction and phage Abs selection

Total RNA was purified from PBMCs using RNeasy mini kit (Qiagen GmbH, Germany). CDNA synthesis was performed using Verso cDNA synthesis kit (Thermoscientific, USA) and used as a template for Abs variable region coding fragments amplification, as detailed in Supplementary. Briefly, heavy and light Ig variable domains (V_H_ and V_L_) were amplified, using specific primer set (supplementary Table 1). The VH and VL used in PCR overlap extension reaction, resulted in scFv repertoire cloned into pCC16 phagemid vector^22^ using *Nco*I/*Not*I. Total of 9.2e6 independent clones obtained, representing the library complexity.

Panning performed against huFc-RBD directly absorbed to polystyrene plates and against biotinylated-huFc-RBD (biotinylation performed using a commercial kit: EZ-Link sulfo-NHS-biotin, Pierce-Thermoscientific, USA) attached to streptavidin coated magnetic beads (Dynabeads Invitrogen, USA). All routine phage display techniques, performed as described ^23^. Screen of specific binders, perform using phage ELISA against huFc-RBD Vs huFc-NTD as control.

### scFv individual clones diversity and sequence verification

TAB-RI_For(CCATGATTACGCCAAGCTTTGGAGCC)andCBD-AS_Rev (GAATTCAACCTTCAAATTGCC) phagemid specific primers used for colony PCR and sequence analysis of scFv Ab individual clones. Colony PCR product, were analyzed on 1.5 % agarose gel, to confirm the intact of the scFv. Restriction fragment size polymorphism (RFLP), performed using MvaI (FastDigest #FD0554; Thermo Scientific, USA) to evaluate sequence variability of scFv individual clones. Following colony PCR, 5µL of the PCR products were taken directly for restriction with 0.5 µL MvaI and 1 µL buffer x10 (provided by the manufacturer) in a 10 µL reaction volume. Restriction was conducted for one hour at 37 °C, and the entire reaction mix was then resolved on 3% agarose gel. Nucleic acid sequence analysis of individual scFv fragments, performed to the colony PCR product, using SeqStudio Genetic Analyzer (Applied Biosystems, USA).

### Expression of SARS-CoV-2 spike recombinant protein

Mammalian cell codon optimized nucleic sequence, coding for SARS-CoV-2 spike glycoprotein (GenPept: QHD43416 ORF), used to design pcDNA3.1+ based expression plasmids, mediating recombinant expression of the entire spike glycoprotein, receptor binding domain (RBD), N-terminal domain (NTD), S1 and S2. Stabilized soluble version of the spike protein (amino acids 1-1207) designed essentially as described earlier ^24, 25^. C-terminal his-tag as well as streptag, included in all constructs in order to facilitate protein purification.

In addition, huFc-RBD fused protein was expressed using previously designed Fc-fused protein expression vector ^26^, giving rise to a protein comprising a homodimer of two RBD moieties (amino acids 318-542, GenPept: QHD43416 ORF) fused to a dimer of IgG1 human Fc (huFc). Expression performed using ExpiCHO^™^ Expression system (ThermoFisher scientific, Gibco^™^).

### Production of scFv-Fc Antibodies

Phagemid DNA of the desired clones were isolated using QIAprep spin Miniprep kit (Qiagen, GmbH, Hilden, Germany), and the entire scFv was cloned into a pcDNA3.1+ based expression vector that was modified, providing the scFv with the human (IgG1) CH2-CH3 Fc fragments, resulting in scFv-Fc antibody format. ScFv-Fc were expressed using ExpiCHO^™^ Expression system (ThermoFisher scientific, Gibco^™^) and purified on HiTrap Protein-A column (GE healthcare, UK).

### ELISA

Routine ELISA protocol applied essentially as described^27^. Coating performed using 2 μg/ml protein. For phage ELISA, HRP-conjugated anti-M13 antibody (Sino Biologicals, USA) was used following detection with TMB substrate (Millipore, USA). ELISA of both sera and recombinant scFv-Fc human antibodies applied with AP-conjugated anti-human IgG (Jackson ImmunoResearch, USA) following detection using PNPP substrate (Sigma, Israel).

### Biolayer interferometry for affinity measurements and epitope binning

Binding studies were carried out using the Octet system (ForteBio, USA, Version 8.1, 2015) that measures biolayer interferometry (BLI). All steps were performed at 30°C with shaking at 1500 rpm in a black 96-well plate containing 200 μl solution in each well. Streptavidin-coated biosensors were loaded with biotinylated antibody (1-5 µg/ml) to reach 0.7-1 nm wavelength shift followed by a wash. The sensors were then reacted for 300 seconds with increasing concentrations of RBD (association phase) and then transferred to buffer-containing wells for another 600 seconds (dissociation phase). Binding and dissociation were measured as changes over time in light interference after subtraction of parallel measurements from unloaded biosensors. Sensorgrams were fitted with a 1:1 binding model using the Octet data analysis software 8.1 (Fortebio, USA, 2015), and the presented values are an average of several repeated measurements. For the binning experiments of antibodies pairs, antibody-loaded sensors were incubated with a fixed RBD concentration (300 nM), washed and incubated with the non-labeled antibody counterpart.

### Cells

Vero E6 (ATCC® CRL-1586^™^) were obtained from the American Type Culture Collection (Summit Pharmaceuticals International, Japan). Cells were used and maintained in Dulbecco’s modified Eagle’s medium (DMEM) supplemented with 10% fetal bovine serum (FBS), MEM non-essential amino acids, 2 nM L-Glutamine, 100 Units/ml Penicilin, 0.1 mg/ml streptomycin and 12.5 Units/ml Nystatin (Biological Industries, Israel). Cells were cultured at 37°C, 5% CO_2_ at 95% air atmosphere.

### Plaque reduction neutralization test (PRNT)

SARS-CoV-2 (GISAID accession EPI_ISL_406862) was kindly provided by Bundeswehr Institute of Microbiology, Munich, Germany. Stocks were prepared by infection of Vero E6 cells for two days. When CPE was observed, media were collected, clarified by centrifugation, aliquoted and stored at -80°C. Titer of stock was determined by plaque assay using Vero E6 cells. Handling and working with SARS-CoV-2 was conducted in BL3 facility in accordance with the biosafety guidelines of the IIBR.

Plaque reduction neutralization test (PRNT), performed as described earlier ^28^. Vero E6 cells were seeded overnight (as detailed above) at a density of 0.5e6 cells/well in 12-well plates. Antibody samples were 2-fold serially diluted (ranging from 100 to 0.78 μg/ml) in 400 μl of MEM supplemented with 2% FBS, MEM non-essential amino acids, 2 nM L-Glutamine, 100 Units/ml Penicilin, 0.1 mg/ml streptomycin and 12.5 Units/ml Nystatin (Biological Industries, Israel). 400 μl containing 300 PFU/ml of SARS-CoV-2 virus were then added to the mAb solution supplemented with 0.25% guinea pig complement sera (Sigma, Israel) and the mixture incubated at 37°C, 5% CO_2_ for 1 hour. Monolayers were then washed once with DMEM w/o FBS and 200 μl of each mAb-virus mixture was added in duplicates to the cells for 1 hour. Virus mixture w/o mAb was used as control. 2 ml overlay [MEM containing 2% FBS and 0.4% tragacanth (Sigma, Israel)] were added to each well and plates were further incubated at 37°C, 5% CO_2_ for 48 hours. The number of plaques in each well was determined following media aspiration, cells fixation and staining with 1 ml of crystal violet (Biological Industries, Israel). NT_50_ was defined as mAb concentration at which the plaque number was reduced by 50%, compared to plaque number of the control (in the absence of Ab).

## Results and Discussion

### Isolation of anti-SARS-CoV-2 antibodies

Several blood samples, either from COVID-19 convalescence or from patients with severe ongoing disease, were evaluated for titers of RBD binding and viral neutralization activity. Two blood samples, exhibiting the highest neutralizing ability (NT_50_ >5000), demonstrating significant binding to both S1 subunit [DIL_50_ (half-dilution value) of 494 and 473] and RBD (DIL_50_ of 252 and 226; Fig. 1a), were subsequently selected for antibody library generation.

**Figure 1.**
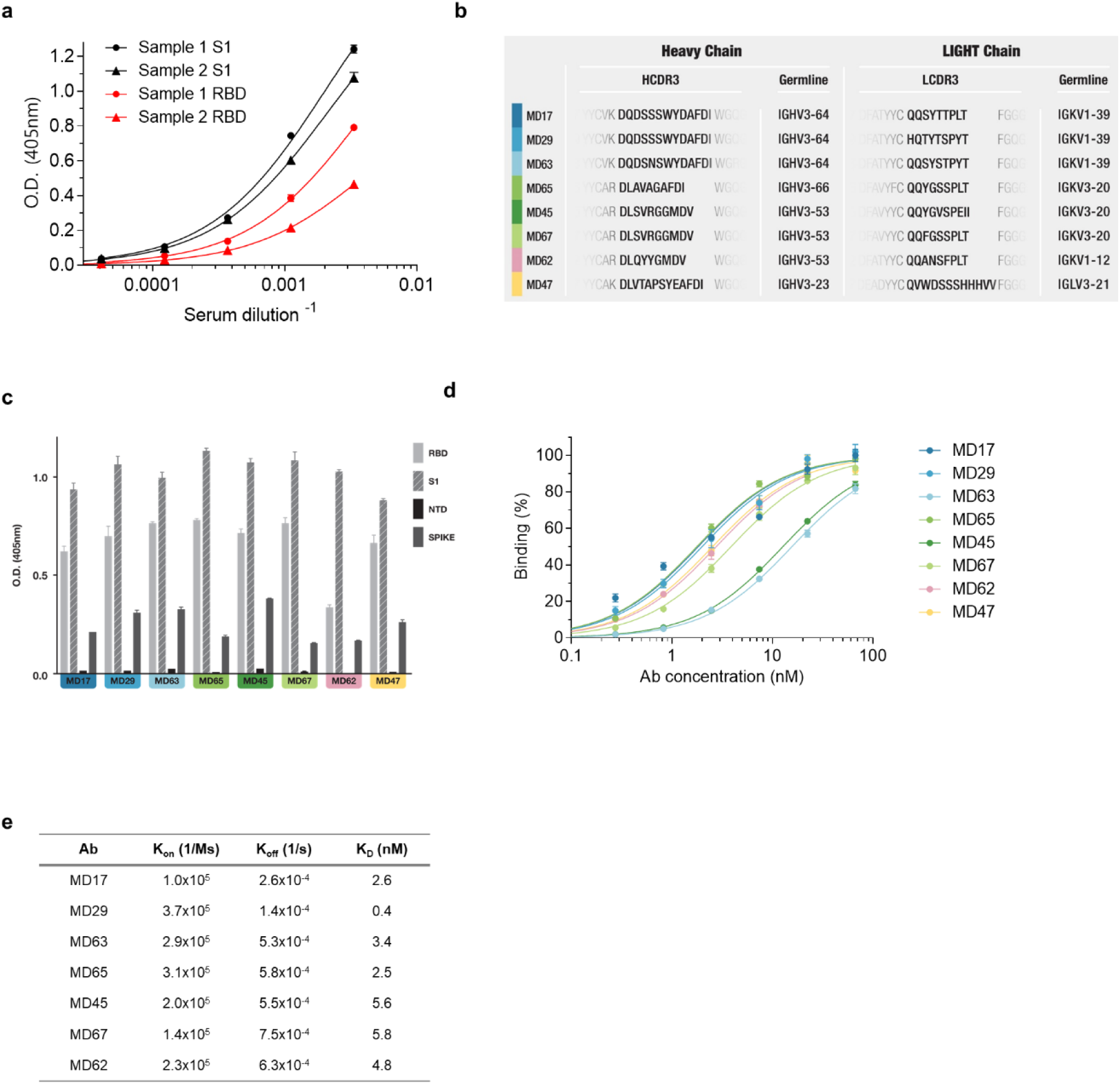
Characterization of the novel human anti-SARS-CoV-2 antibodies. **a**. Binding curves of polyclonal antibodies in serially diluted serum samples of COVID-19 patients obtained by ELISA using S1 or RBD as coating antigen. **b**. Amino acid sequences of the HCDR3 and LCDR3 of the selected antibodies and their respective germ line genes. **c**. Specificity of the selected antibodies determined by ELISA against the indicated SARS-CoV-2 proteins. **d**. Reactivity profile of antibodies determined by ELISA, using S1 as the coating antigen. Data is presented as binding percent of Bmax for each antibody. **e**. Binding characteristics of the monoclonal antibodies determined using biolayer interferometry. All antibodies (except MD47) were biotinylated, immobilized to the sensor and interacted with increasing amounts of RBD. Binding kinetics were fitted using the 1:1 binding model. The values shown represent average ± SEM.

A phage display (PD) single chain Fv (scFv) library, representing approx. 10 million distinct antibodies, was constructed. With the objective of isolating neutralizing Abs, three consecutive enrichment steps of panning were performed against both S1 and RBD. Resulting clones were tested for their ability to bind S1, the positive once were expressed as full-length antibodies (in a scFv-Fc format) for further analysis. Subsequently, eight RBD-specific antibodies carrying unique sequences were selected (Fig 1b).

Binding specificity assays of these eight antibodies confirmed their specificity to the spike protein, the S1 subunit and the RBD, while no reactivity was observed against the Spike N terminal domain (NTD) of the spike protein (Fig. 1c). Evaluation of the Abs affinity toward S1 by ELISA evidenced apparent K_D_ values of 1.8-3.8 nM of six of the antibodies, and a K_D_ values of 12.7 and 15.6 nM of MD45 and MD63, respectively (Fig. 1d). To further characterize these antibodies, biolayer interferometry (BLI) measurements of the antibodies affinity specifically toward RBD was conducted, revealing K_D_ values of 0.4 to 5.8 nM for all antibodies, with MD29 showing the highest affinity and MD67 showing the lowest affinity (Fig. 1e). It should be noted that the biotinylation of MD47 was found to significantly hinder its ability to bind RBD and therefore was not evaluated in this format. Additional sequence analysis by IgBlast^18^ (Fig 1b) revealed that antibodies MD17, MD29 and MD63 share common germ line origin of both their VH and VK (IGHV3-64 and IGKV1-39, respectively). Similarly, MD45 and MD67, share common VH and VK germ lines (IGHV3-53 and IGKV3-20). The VH of MD62, is similar to the one of MD45 and MD67, accompanied by a unique VK (IGKV1-12), while MD65 shared the same VK with MD45 and MD67, accompanied by a unique VH (IGHV3-66). MD47 originated from unique VH (IGHV3-23) and was the only mAb, carrying VL (IGLV3-21).

### Classification of antibody epitopes

SARS-CoV-2 spike RBD is known to mediate the binding of the human ACE2 receptor and thus, this domain is considered as main target for neutralizing mAbs. However, direct blocking of the RBD-ACE2 interaction is not the exclusive modality by which neutralizing antibodies can exert their effect^21^. Consequently, the selected mAbs, were classified on the basis of their epitope specificity determined by BLI epitope binning. In this assay, each individual antibody was biotinylated, immobilized to a streptavidin sensor, loaded with RBD and then challenged with each of the other antibodies. Simultaneous binding of the second antibody to RBD induces a wavelength shift in the interference pattern, which indicates that the two antibodies bind to non-overlapping epitopes^19^. Conversely, if the two antibodies bind the same or partially-overlapping epitope on RBD, no or very low wavelength shift, respectively, is induced. As a representative example, sensograms of the various antibody interactions with a pre-complexed MD65-RBD is shown (Fig. 2a). Antibody MD65 used as a negative control, and did not elicit any wavelength shift, as expected. In contrast, antibodies MD29, MD47, MD62 and MD63 induced a marked wavelength shift indicating that they could bind to RBD simultaneously with MD65. The analysis revealed that antibodies MD45 and MD67 could not bind to RBD in the presence of MD65, indicating that these three antibodies target the same epitope. Analysis was then performed for the next seven antibodies and the ability of each pair to simultaneously bind RBD was determined (Fig. 2b). Results ranged from full to no competition and enabled the classification of the mAbs into 4 groups recognizing distinct epitopes (Fig 2c): I (MD17, MD29 and MD63); II (MD45, MD65 and MD67); III (MD62) and IV (MD47).

**Figure 2.**
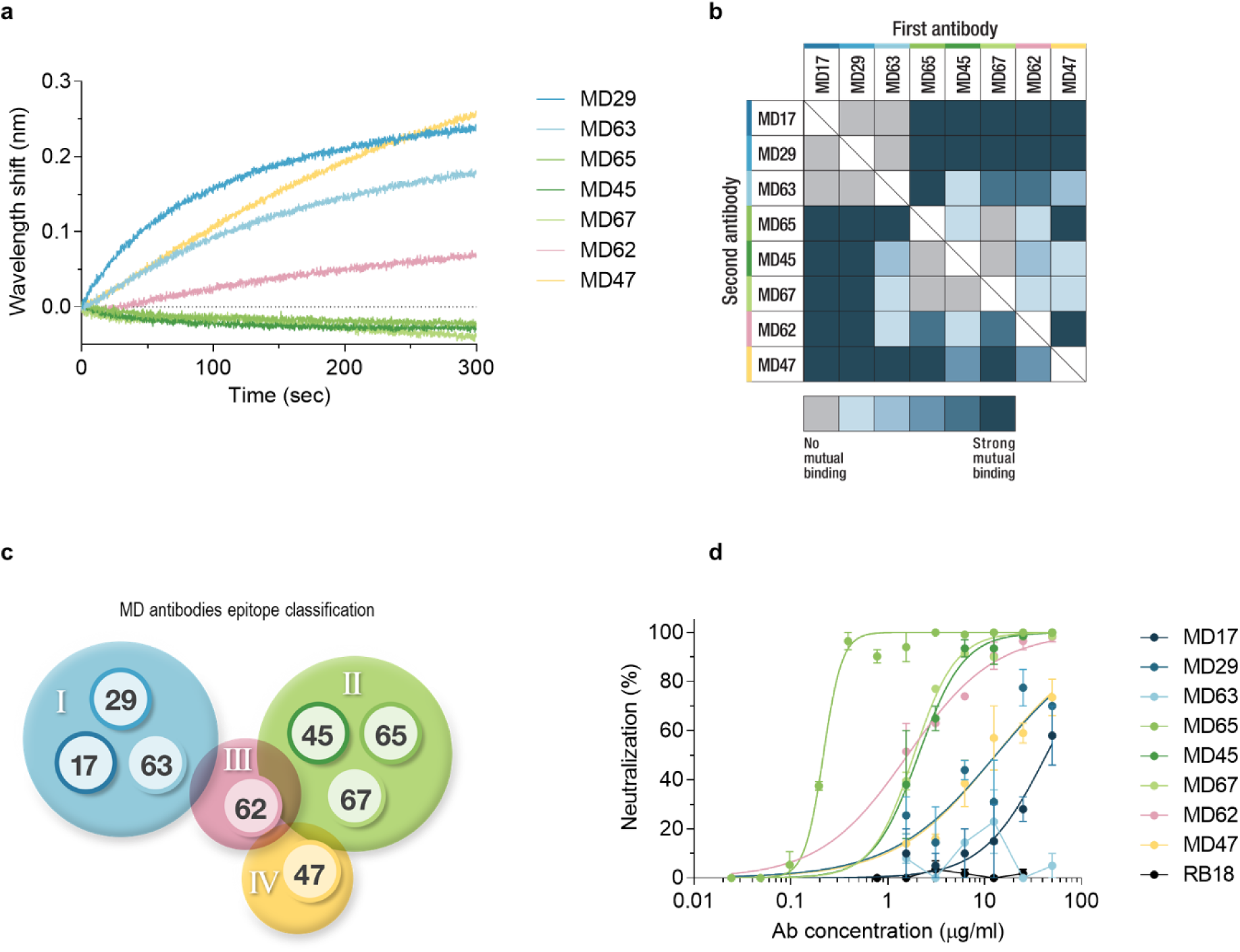
Epitope binning and SARS-CoV-2 neutralization. **a**. Biolayer interferometry was applied for the epitope binning experiments. Representative assay results are shown for MD65 mAb. The purified antibody was biotinylated, immobilized on streptavidin sensor and saturated with RBD. The complex was then incubated for 300 sec with each one of the indicated antibodies. Time 0 represents the binding to the MD65-RBD complex. **b**. Complete epitope binning of the eight selected MD monoclonal antibodies. Binding was evaluated by the ability of each pair of antibodies to simultaneously bind RBD, using biolayer interferometry. **c**. Four non-competing RBD binding epitopes were identified and accordingly classified into four groups: I (blue), II (green), III (pink) and IV (yellow). **d**. SARS-CoV-2 *in vitro* neutralization using plaque reduction neutralization test (PRNT). Neutralization potency was determined by the ability of each antibody (at indicated concentrations) to reduce plaques formation; results are expressed as percent inhibition of control without Ab. The values shown represent average ± SEM.

Most notably, the classification of the antibodies on the basis of their specific targeted antigenic epitopes, is strongly supported by their observed sequence similarity, as discussed above. Group I mAbs shared the same V germ lines and were found to target the same epitope. Similarly, mAbs MD45 and MD67 which shared the same V germ lines and the same epitope, were classified as group II. Although MD65 differ in its VH germ line, he was included in this group as well. On the other hand, MD62 sharing the same VH germ line with MD45 and MD67, appears to bind a distinct epitope (III). Finally, MD47 exhibited both a unique sequence and targeted a unique epitope (Fig. 2c).

Recently, the RBD-located epitope recognized by the SARS-CoV specific antibody CR3022, was determined^14,20^. We chose to use this antibody in order to further determine the epitopes recognized by the antibodies described in this report. Therefore, a recombinant in-house version of this antibody was generated (Supplementary data) and used in epitope binning assays, together with the selected set of novel mAbs. CR3022 IgG was immobilized to the BLI sensor, loaded with RBD and further challenged with each of the selected mAbs. Group I mAbs were found to compete with the CR3022 as evidenced by the lack of interaction with the mAb-RBD complex (Supplementary Fig. 1). The remaining five tested mAbs did bind the RBD in the presence of the CR3022 antibody. These results suggest that antibodies MD17, MD29 and MD63 bind to the RBD epitope that spans the RBD residues 369-386, previously defined as the CR3022 epitope^20^.

### SARS-CoV-2 neutralization by the novel human antibodies

The neutralization potency of the antibodies was evaluated by plaque reduction neutralization test (PRNT) using VeroE6 cells infected with the pathogenic SARS-CoV-2. A fixed amount of the virus was incubated with increasing concentrations of each antibody, the mixture was then added to the cells, and the number of plaques was quantified 48 hours later. Antibodies MD45, MD67, MD62 and MD65 displayed the highest neutralization potency, with a neutralization dose needed to inhibit 50% of the plaques (NT_50_) of 2.1, 1.9, 1.6 and 0.22 μg/ml, respectively. MD65 exhibited the highest neutralization capacity amongst the entire set of antibodies (Fig. 2d). Interestingly, antibodies MD17, MD29 and MD63 shared DNA sequence homology indicative of a common germ line and competed for the same epitope, yet only the first two showed neutralizing activity (NT_50_ of 43 and 13 μg/ml, respectively). This discrepancy may be explained by differences in the affinities of the antibodies in this group. A recent report similarly documented that antibody CR3022 (targeting an epitope overlapping with MD17 and MD29, see above) does not neutralize the novel SARS-CoV-2 presumably due to its low binding affinity to the RBD (115 nM)^20^. Additional antibody tested in this study, MD47, which represents a unique epitopic group, also showed neutralizing activity with NT_50_ of 13 μg/ml. Note that irrelevant Ab, RB18 (scFv-Fc) targeting abrin toxin, showed no neutralization against the virus.

To conclude, we report the isolation and characterization of a set of fully human SARS-CoV-2, neutralizing antibodies that target four distinct epitopes on the spike RBD. As neutralizing antibodies are generally known to be useful as post-exposure therapy for viral infection and more specifically for treatment of human corona viral diseases^3, 21^, we suggest that these antibodies might serve as an efficient treatment of COVID-19 patients or for prophylaxis immunization. Furthermore, since these neutralizing antibodies target different epitopes, they can be combined to further improve treatment efficacy and to reduce the risk of the emergence of treatment-escaping viral variants.

## Acknowledgments

We thank Dr. Itzchak Levy and Dr. Asaf Biber from the Sheba Medical Center and Dr. Hagar Mizrahi, Dr. Hiba Zayyad and Dr. Moshe Matan from the Baruch Padeh Medical Center, for kindly providing the blood samples collected from the COVID-19 patients. We wish to express our gratitude to our colleagues Dr. Ofer Cohen, Dr. Emanuelle Mamroud, Moshe Mantzur, Moshe Aftalion, Noa Caspi, Yentl Evgy, Hila Cohen, Dr. Liat Bar-On, Dr. Ronit Aloni-Grinstein, David Gur, Sarit Sterenberg and Tamar Aminov for fruitful discussions and support. We acknowledge the IIBR administrative personnel for their commitment to the project and Dalit Brener for graphical design.

## Author Contributions

R.R., O.M., T.N-P. and E.M. conceived and designed the experiments and wrote the manuscript. T.N-P., E.M., R.A., A.M., Y.L., A.B-K., A.Z., H.T., Y.Y-R., M.I., E.E., H.A., S.M., T.C., S.W., E.P., O.R., N.P., T.I., O.M. and R.R. designed and performed the experiments and analyzed the data. S.Y. and S.S added fruitful discussions. R.R. and O.M. supervised the project.

