## Supplementary data for "Tiger team: a panel of human neutralizing mAbs targeting SARS-CoV-2 spike at multiple epitopes"

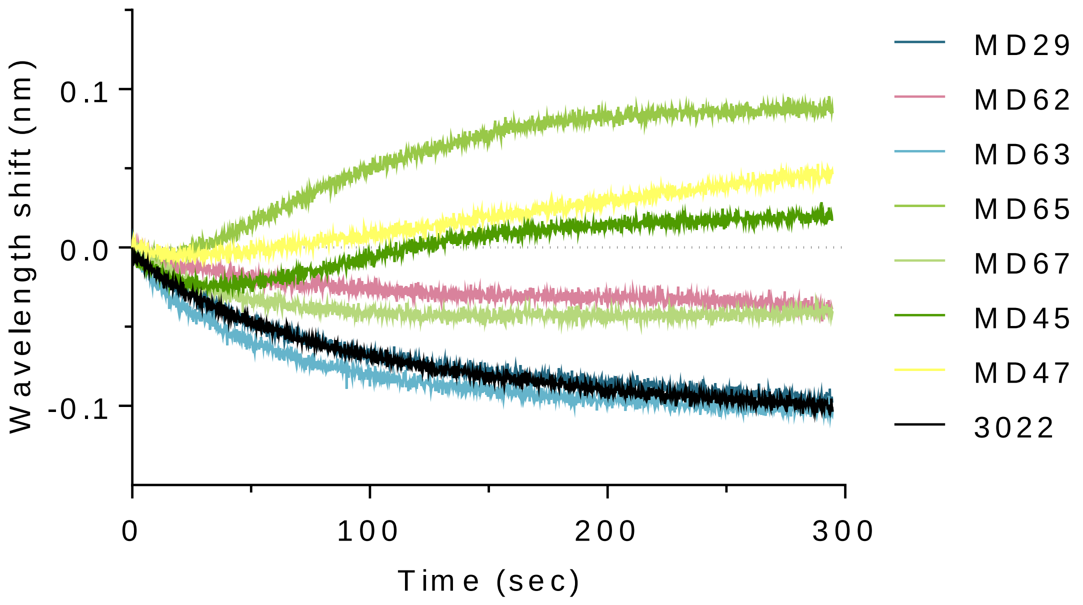


**Supplementary Figure 1. Epitope binning of the selected antibodies versus CR3022**. Biolayer interferometry was applied for this epitope binning experiment. Immobilized antibody CR3022 was saturated with RBD and then the complex was incubated for 300 sec with each one of the indicated antibodies. Time 0 represents the binding to the CR3022-RBD complex.

Supplementary Table 1. Primer set for human phage display (PD) library construction

| Primer name | Primer Sequence |
|---|---|
| **Heavy chain variable region amplification primers** | |
| Hu -VH-PD_For1 | ctttctatgcggcccagccggccatggcc CAGGTBCAGCTKGTRCARTCTGG |
| Hu -VH-PD_For2 | ctttctatgcggcccagccggccatggcc CARATGCAGCTGGTGCAGTCTGG |
| Hu -VH-PD_For3 | ctttctatgcggcccagccggccatggcc GARGTSCAGCTGGTRCAGTCTGG |
| Hu -VH-PD_For4 | ctttctatgcggcccagccggccatggcc CAGATCACCTTGAAGGAGTCTGG |
| Hu -VH-PD_For5 | ctttctatgcggcccagccggccatggcc CAGGTCACCTTGAGGGAGTCTGG |
| Hu -VH-PD_For6 | ctttctatgcggcccagccggccatggcc CAGGTCACCTTGAAGGAGTCTGG |
| Hu -VH-PD_For7 | ctttctatgcggcccagccggccatggcc GARGTRCARCTGGTGGAGTCYGG |
| Hu -VH-PD_For8 | ctttctatgcggcccagccggccatggcc CAGGTGCAGCTGGTGGAGTCTGG |
| Hu -VH-PD_For9 | Ctttctatgcggcccagccggccatggcc GAGGTGCAGCTGTTGGAGTCTGG |
| Hu -VH-PD_For10 | Ctttctatgcggcccagccggccatggcc GAGGTGCAGCTGGTGGAGWCTG |
| Hu -VH-PD_For11 | Ctttctatgcggcccagccggccatggcc CAGGTGCARCTGCAGGAGTCGGG |
| Hu -VH-PD_For12 | Ctttctatgcggcccagccggccatggcc CAGCTGCAGCTGCAGGAGTCSGG |
| Hu -VH-PD_For13 | Ctttctatgcggcccagccggccatggcc CAGGTACAGCTGCAGCAGTCAGG |
| Hu -JH-PD_Rev1 | accaccaccaccggatcctcctcctcctgctgagcc TGAGGAGACRGTGACCAGGG |
| Hu -JH-PD_Rev2 | accaccaccaccggatcctcctcctcctgctgagcc TGAAGAGACGGTGACCATTGTCC |
| Hu -JH-PD_Rev3 | accaccaccaccggatcctcctcctcctgctgagcc TGAGGAGACGGTGACCGTGGTCC |
| **Kappa chain variable region amplification primers** | |
| Hu-Vk-PD_For1 | ggatccggtggtggtggttctggcggcggcggctcc RHCATCYRGWTGACCCAGTC |
| Hu-Vk-PD_For2 | ggatccggtggtggtggttctggcggcggcggctcc GAYRTYGTGATGACYCAGWC |
| Hu-Vk-PD_For3 | ggatccggtggtggtggttctggcggcggcggctcc GAAATWGTRWTGACRCAGTC |
| Hu-Vk-PD_For4 | ggatccggtggtggtggttctggcggcggcggctcc GAAACGACACTCACGCAGTC |
| Hu-Vk-PD_For5 | ggatccggtggtggtggttctggcggcggcggctcc GAWRTTGTGMTGACWCAGTC |
| Hu-JK-PD_Rev1 | gataccggtgtatttgcgccacctgcggccgc TTTGATHTCCACYTTGGTCC |
| Hu-JK-PD_Rev2 | gataccggtgtatttgcgccacctgcggccgc TTTGATCTCCAGCTTGGTCC |
| Hu-JK PD_Rev3 | gataccggtgtatttgcgccacctgcggccgc TTTAATCTCCAGTCGTGTCC |
| **Lambda chain variable region amplification primers** | |
| Hu-VL-PD _For1 | ggatccggtggtggtggttctggcggcggcggctcc CAGTCTGTSBTGACKCAGCC |
| Hu-VL-PD _For2 | ggatccggtggtggtggttctggcggcggcggctcc CAGTCTGCCCTGACTCAGCC |
| Hu-VL-PD _For3 | ggatccggtggtggtggttctggcggcggcggctcc TCYTMTGWGCTGACWCAGCC |
| Hu-VL-PD _For4 | ggatccggtggtggtggttctggcggcggcggctcc TCCTATGAGCTGAYHCAGSWVC |
| Hu-VL-PD _For5 | ggatccggtggtggtggttctggcggcggcggctcc CAGSYTGTGCTGACTCAAYC |
| Hu-VL-PD _For6 | ggatccggtggtggtggttctggcggcggcggctcc AATTTTATGCTGACTCAGCC |
| Hu-VL-PD _For7 | ggatccggtggtggtggttctggcggcggcggctcc CAGRCTGTGGTGACYCAGG |
| Hu-VL-PD _For8 | ggatccggtggtggtggttctggcggcggcggctcc CWGSCWGKGCTGACTCAGCC |
| Hu-JL-PD_Rev1 | gataccggtgtatttgcgccacctgcggccgc TAGGACGGTSACCTTSGTCCC |
| Hu-JL-PD_Rev2 | gataccggtgtatttgcgccacctgcggccgc TAGGACGATCAGCTGGGTTCC |
| Hu-JL-PD_Rev3 | gataccggtgtatttgcgccacctgcggccgc TAGGACGGTCAGCTCSGTCCC |
| Hu-JL-PD_Rev4 | gataccggtgtatttgcgccacctgcggccgc TAGGACGGTCASCTKGGTKCC |
| **Single chain assembly primers** | |
| ASS1_For | CTTTCTATGCGGCCCAGC |

| | |
|---|---|
| ASS1_Rev | GATACCGGTGTATTTGCGCC |
| **Single chain_pCC16 amplification primers** | |
| TAB-RI-For | CCATGATTACGCCAAGCTTTGGAGCC |
| CBD-As-Rev | GAATTCAACCTTCAAATTGCC |

All PCR reactions contained Advantage 2 DNA polymerase mix and reaction buffer (Clontech, USA), 10 mM dNTPs (Promega, USA) and 0.5 µM of each primer [Desalted oligonucleotides (Sigma, Israel)] in total of 25 µl volume.

**Phage-display library construction**

Heavy and light Ig variable domains (VH and VL) amplification was performed directly from cDNA, using V/J specific set of primers (indicated by upper case in Table S1). Amplification performed separately for each primer pair (1 min 95°C; 30 cycles of: 20 s, 94°C; 30 s, 55°C; 1 min, 72°C; 5 min, 72°C). PCR products were directly used as a template for a second amplification (1 min 95°C; 20 cycles of: 20 s, 94°C; 30 s, 57°C; 1 min, 72°C; 5 min, 72°C), using the above listed primers, adding flanking sequences (indicated by lower case) introducing restriction recognition sites for subsequent cloning into pCC16 phagemid vector. Following gel purification of the variable fragments [using QIAquick Gel extraction kit (Qiage, Germany)] assembly PCR performed for 200 ng of VH and Vκ/Vλ products (in a 1:1 ratio). First 10 assembly cycles were performed without primers addition (95°C, 1 min; 10 cycles of: 95°C, 15 s; 60°C, 30 s; 72°C, 30 s; 72°C, 5 min). Next, products were diluted 1:10 in fresh reaction mixture including the ASS1 primer pair for an additional 30 cycles (95°C, 1 min; 30 cycles of: 95°C, 15 s; 65°C, 20 s; 72°C, 1 min; 72°C, 5 min).

For PD library construction, pCC16 plasmid and assembled scFvs were digested with 5 U each of *Nco*I and *Not*I restriction enzymes (FastDigest, Thermo Scientific, USA), in a 50 µL reaction volume, 2 hrs. at 37°C. The resulted pCC16 vector was purified from 1% agarose gel using the QIAquick Gel extraction kit, and the resulted scFv inserts were purified using the QIAquick PCR purification kit (Qiagen, Germany). The scFv was than cloned into 200 ng of the pCC16 vector (5:1 ratio; insert: vector), in a 15 µL reaction volume, using 6 U of ligase (T4 HC ligase; Thermo Scientific, USA), with 2 hr incubation at RT and further overnight incubation at 4°C, followed by inactivation at 70°C for 5 min. Each ligation reaction was transformed to 30 µL of E. coli TG1 electrocompetent cells (Lucigen, USA), and a total of 48 cloning reactions were performed. The transformed bacteria, containing the final scFv libraries, were plated on YPD agar (BD, USA) supplemented with 100 µg/mL ampicillin and 100 mM glucose and, after an overnight culture at 30°C, were harvested, aliquoted and stored at -80°C.
